# Chemotaxonomy supports morphology in the identification of *Vepris hemp* (Rutaceae) a new species of Critically Endangered deciduous forest shrub from Usambara Mts, Tanzania

**DOI:** 10.1101/2022.08.27.505527

**Authors:** Moses Langat, Andreas Hemp, Martin Cheek

## Abstract

*Hemp* 7152, a sterile herbarium plot voucher of a shrub from a rare type of deciduous forest in the Usambara Mts, Tanzania was tentatively identified using morphology as a new species of *Vepris* (Rutaceae). To gain further support for its placement its chemistry was investigated. The compounds isolated from *Hemp 7152* were four quinoline alkaloids, kokusaginine (**1**), *N*-methylplaty-desminium ion (**9**), ribalinium ion (**10**), and isoplatydesmine (**11**), and seven acridone alkaloids, arborinine (**2**) 1,2,3-trimethoxy-*N*-methylacridone (**3**), 1,2,3,5-tetramethoxy-*N*-methylacridone (**4**), 1,3-dimethoxy-*N*-methylacridone (**5**) and toddaliopsis A (**6**), evoxanthine (**7**) and tecleanthine (**8**). In addition, lupeol and ferulic acid were isolated from this plant. The combination of quinoline and acridone alkaloids is restricted to the *Rutaceae* family, confirming beyond reasonable doubt the placement of this material in the Rutaceae. Within Rutaceae in tropical Africa, only the genus *Vepris* is unarmed, with trifoliolate leaves. Using an identification key, and herbarium specimen matching, *Hemp* 7152 was morphologically placed as close to *Vepris uguenensis*, sharing xerophytic characters unusual in the genus. The species are geographically close, occurring in adjoining mountains in northern Tanzania. However, *Vepris uguenensis* contains 13 alkaloids which are not present in *Hemp* 7152, nor in any other species of *Vepris* that has been studied, supporting species recognition for *Hemp* 7152 which is formally named as *Vepris hemp*, morphologically characterised, illustrated and assessed as Critically Endangered using the IUCN 2012 standard. The new species appears restricted to an almost extinct type of deciduous forest, characterised in this paper.

## INTRODUCTION

As part of a series of studies of the chemistry of *Vepris* (Rutaceae) led by the first author, material from a sterile, morphologically distinct and unmatched *Vepris* taxon collected by the second author was investigated. In this paper we present the chemical results supporting its placement in Rutaceae, specifically *Vepris*, compare the taxon morphologically within *Vepris*, test the hypothesis that this taxon is new to science based on the combination of chemical and morphological data, and formally name it as *Vepris hemp* Cheek & Langat. We also present data on its ecology in a rare, unprotected, surviving fragment of an unusual type of deciduous forest in the Usambara Mts of Tanzania, East Africa.

Since 1996 the second author has led the placement and enumeration of 2500 vegetation plots in Ethiopia, Kenya, and the greatest number, in Tanzania (*Hemp, 2012*, see also Methods below). The plot network is aimed at analysing diversity patterns along climatic and land-use gradients in East Africa. Specimens from the plots, both fertile and sterile, are pre-identified at the herbaria of the field station of the KiLi-Project (https://www.kilimanjaro.biozentrum.uni-wuerzburg.de/) on Kilimanjaro and the National Herbarium of Tanzania (NHT). Those requiring further identification are taken mainly to the Kew herbarium (K) where the reference specimens are more comprehensive and taxonomic specialists can support expert identifications. From time-to-time new species to science are uncovered, e.g. *Garcinia tanzaniensis* Verdc. (Clusiaceae, *Verdcourt 2007*), *Lellingeria paucipinnata* Parris (Grammitidaceae, *Parris, 2002*), *Chlorophytum rhizopendulum* Bjorå & Hemp (Anthericaceae, *Sletten Bjorå et al., 2008*), *Crotalaria arrecta* Hemp & Polhill (Leguminosae, *Hemp & Polhill, 2009*), *Asplenium arcumontanum* Hemp & N.R.Crouch (Aspleniaceae, *Hemp & Crouch, 2018*), *Pimpinella silvicola* (*Hemp, 2015*) and *Rhipidoglossum monteparense* (Orchidaceae, *Cribb & Hemp, 2022*).

In 2019 the second author was seeking to identify *Hemp* 7152, a sterile plot voucher from an unarmed shrub with alternate, astipulate, trifoliolate, gland-dotted leaves which was tentatively identified as the genus *Vepris* (Rutaceae). The third author, a taxonomic specialist in this genus, concurred with the generic identification, and established that it matched no other known species of the genus in tropical Africa (see results). The site of collection, the West Usambara Mts, is a renowned centre of diversity with numerous point endemics, which are steadily increased by the addition of new species to science. It was assumed that *Hemp* 7152 represented a new species to science. However, the standard convention in angiosperm plant taxonomy is that species are not described as new to science unless flowers and/or fruits are present (*Cheek et al., 2020*). Sterile specimens, lacking such structures, are usually set aside until this deficiency can be addressed. The logic for this convention is partly because flowers and or fruits are often essential to confirm generic placement of a specimen. In the case of *Vepris*, the genus is so distinct vegetatively that the possibility of confusing it with any other genus in continental tropical Africa is remote. Nonetheless, firm evidence of placement in Rutaceae would give a concrete basis for description of the new species. Therefore, the first author, a plant chemist was approached to characterise the secondary compounds in the specimen, since Rutaceae are characterised by a suite of compounds, particularly acridones and quinolines which in combination are unique to the family (*Basa & Tripathy, 1984; da Silva, Gottlieb & Ehrendorfer, 1988; Imbenzi et al., 2014; Michael, 2007; Michael, 2017*; *Ombito et al., 2021; Sackett, 2000*). The outcome provided this data (see results), supporting the formal description of the new species, which is necessary to support formal IUCN conservation assessment, needed to improve the possibility of protecting what appears to be a rare and threatened species. Accordingly, the specimen is chemically characterised, formally described morphologically, and named in this paper.

*Vepris* Comm. ex A. Juss. (Rutaceae-Toddalieae), is a genus with 91 accepted species, 18 in Madagascar and the Comores and 73 in Continental Africa with one species extending to Arabia and another endemic to India (POWO, continuously updated). The genus was last revised for tropical Africa by *Verdoorn (1926)*. Founded on the Flore du Cameroun account of *Letouzey* (*1963*), eight new species were recently described from Cameroon (*Onana & Chevillotte 2015; Cheek, Gosline & Onana, 2018; Onana, Chevillotte, Cheek, 2019; Cheek & Onana, 2021; Cheek, Hatt & Onana, 2022*), taking the total in Cameroon to 24 species, the highest number for any country globally. The greatest concentration of *Vepris* species in Cameroon is within the Cross-Sanaga Interval (*Cheek et al., 2001*) with 15 species of *Vepris* of which nine are endemic to the Interval. The Cross-Sanaga has the highest species and generic diversity per degree square in tropical Africa (*Barthlott, Lauer & Placke, 1996; Dagallier et al., 2020*) including endemic genera such as *Medusandra* Brenan (Peridiscaceae, *Breteler, Bakker & Jongkind, 2015; Soltis et al., 2007*).

Multiple Cameroonian *Vepris* species are threatened (*Onana & Cheek, 2011*) and one species is globally extinct (*Cheek, Gosline & Onana, 2018*).

In other parts of Africa species are also highly threatened, e.g., the Critically Endangered *Vepris laurifolia* (Hutch. & Dalziel) O. Lachenaud & Onana of Guinea-Ivory Coast (formerly *V. felicis* Breteler, *Cheek, 2017; Lachenaud & Onana, 2021*), and in Tanzania, the still unpublished *Vepris* sp. A of FTEA is considered extinct (*Cheek & Luke, 2022*).

In mainland tropical Africa, *Vepris* are readily characterised. Among other Rutaceae they differ because they have digitately (1–)3(–5)-foliolate leaves, and unarmed stems. The other genera are pinnately compound and often spiny. *Vepris* species are evergreen shrubs and trees of tropical lowland evergreen forest, extending into cloud forests and a few into drier forest and woodland. *Vepris* species are often indicators of good quality evergreen forest since they are not pioneers. New species are steadily coming to light (*Cheek & Luke, 2022; Langat, Kami & Cheek, 2022*).

*Vepris* species in Africa extend from the fringes of the Sahara Desert (*Vepris heterophylla* (Engl.) Letouzey) to South Africa, e.g. *Vepris natalensis* (Sond.) Mziray. *Mziray (1992)* subsumed the genera *Araliopsis* Engl., *Diphasia* Pierre, *Diphasiopsis* Mendonça, *Oricia* Pierre, *Oriciopsis* Engl., *Teclea* Delile, and *Toddaliopsis* Engl. into *Vepris*, although several species were only formally transferred subsequently (e.g., *Harris, 2000; Gereau, 2001; Cheek, Oben & Heller, 2009; Onana & Chevillotte, 2015*). Mziray’s conclusions were for the most part confirmed by the molecular phylogenetic studies of *Morton (2017)* but Morton’s sampling was limited, identifications appeared problematic (several species appear simultaneously in different parts of the phylogenetic trees) and more molecular work is desirable. Morton studied about 14 taxa of *Vepris*, all from eastern Africa. Recently *Appelhans & Wen (2020)* focussing on Rutaceae of Madagascar, have found that the genus *Ivodea* Capuron is sister to *Vepris* and that a Malagasy *Vepris* is sister to those of Africa. However, the vast majority of the African species including all those of West and Congolian Africa, remain molecularly unsampled leaving the possibility open of changes to the topology of the phylogenetic tree if this is addressed.

Characteristics of some of the formerly recognised genera are useful today in grouping species. The “araliopsoid” species have firm, subglobose, 4-locular fruit, syncarpous with 4 external grooves; the “oriciopsoid” soft, fleshy 4-locular syncarpous fruit; “oricioid” species are 4-locular and apocarpous in fruit; the fruits of “diphasioid” species are laterally compressed in one plane, bilocular and bilobed at the apex; diphasiopsoid” species have two locules entirely free from each other; while “tecleoid” species are unilocular in fruit and 1-seeded, lacking external lobes or grooves. There is limited support for these groupings in Morton’s study,

Due to the essential oils distributed in their leaves, and the alkaloids and terpenoids distributed in their roots, bark and leaves, species of *Vepris* often have medicinal and other values (*Burkill, 1997*). Burkill details the uses, essential oils and alkaloids known from five species in west Africa: *Vepris hiernii* Gereau (as *Diphasia klaineana* Pierre), *Vepris suaveolens* (Engl.) Mziray (as *Teclea suaveolens* Engl.), *Vepris afzelii* (Engl.) Mziray (as *Teclea afzelii* Engl.), *Vepris heterophylla* (Engl.) Letouzey (as *Teclea sudanica* A. Chev.) and *Vepris verdoorniana* (Exell & Mendonça) Mziray (as *Teclea verdoorniana* Exell & Mendonça) (*Burkill, 1997*: 651–653). Research into the characterisation and anti-microbial and anti-malarial applications of alkaloid and limonoid compounds in *Vepris* is active and ongoing (e.g., *Atangana et al., 2017*), although sometimes published under generic names no longer in current use, e.g. *Wansi et al., (2008)*. Applications include as synergists for insecticides (*Langat, 2011*). *Cheplogoi et al*., (*2008*) and *Imbenzi et al*., (*2014*) respectively list 14 and 15 species of *Vepris* that have been studied for such compounds. A review of ethnomedicinal uses, phytochemistry, and pharmacology of the genus *Vepris* was recently published by *Ombito et al*., (*2021*), listing 213 different secondary compounds, mainly alkaloids and furo- and pyroquinolines, isolated from 32 species of the genus, although the identification of several of the species listed needs checking. However, few of these compounds have been screened for any of their potential applications. Recently, *Langat et al*., (*2021*) have published three new acridones new to science and reported multi-layered synergistic anti-microbial activity from *Vepris gossweileri* (I.Verd.) Mziray, recently renamed as *Vepris africana* (Hook.f ex Benth.) Lachenaud & Onana (*Lachenaud & Onana, 2021*).

## MATERIALS & METHODS

### Taxonomy

*The electronic version of this article in Portable Document Format (PDF) will represent a published work according to the International Code of Nomenclature for algae, fungi, and plants (ICN), and hence the new names contained in the electronic version are effectively published under that Code from the electronic edition alone. In addition, new names contained in this work which have been issued with identifiers by IPNI will eventually be made available to the Global Names Index. The IPNI LSIDs can be resolved and the associated information viewed through any standard web browser by appending the LSID contained in this publication to the prefix “http://ipni.org/“. The online version of this work is archived and available from the following digital repositories: PeerJ, PubMed Central, and CLOCKSS*.

Fieldwork in Tanzania resulting in the specimens and observations cited in this paper was conducted with the collaboration and support of the National Herbarium (NHT) at the Tropical Pesticide Research Institute (TPRI) in Arusha and the University of Bayreuth, Germany in 2018 under research permit 2018-300 NA 1996-44 xx (issued 1 June 2018), and the specimens were exported under the Material Transfer Agreement (MTA) between TPRI and the KiLi-Project dated February 2012. Field work was funded by the German Research Foundation (DFG), grant no HE2719/11-3. Duplicates were deposited at NHT, K and UBT.

The taxonomic study is based on herbarium specimens studied by the second two authors at K in 2019 and 2022, and observations of live material in Tanzania made by the second author in 2018. All specimens cited have been seen. The specimen was collected using standard methodology e.g. as in *Cheek & Cable (1997)*. The description was made using the standard of *Langat, Kami & Cheek*, (2022) and terms as in *Beentje & Cheek* (2003). Herbarium citations follow Index Herbariorum (*Thiers et al., continuously updated*), nomenclature follows *Turland et al., (2018)* and binomial authorities follow *IPNI (continuously updated)*. Material of the suspected new species was compared morphologically with material of all other species African *Vepris*, principally at K, but also using material and images from BM, EA, BR, FHO, G, GC, HNG, P and YA. Herbarium material was examined with a Leica Wild M8 dissecting binocular microscope fitted with an eyepiece graticule measuring in units of 0.025 mm at maximum magnification. The drawing was made with the same equipment using a Leica 308700 camera lucida attachment.

### Ecology

Vegetation plots were established following the method of *Braun-Blanquet* (*1964*), which includes information about the vegetation structure and the whole species composition in the different vegetation layers with their cover and frequency. pH was measured in the main root horizon using a WTW pH-meter (pH 330). Two parallel samples were taken and measured in distilled water and a 0.01 M CaCl_2_ solution, respectively.

### Chemistry

The chemistry of the leaves of *Vepris hemp* were analysed using the protocols described in *Langat, Kami & Cheek(2022)*. The FTIR spectra were recorded using a Perkin-Elmer Frontier/Spotlight 200 spectrometer, whereas the 1D and 2D NMR spectra were recorded in CDCl_3_ or CD_3_OD, depending on their solubilities, on a 400 MHz Bruker AVANCE NMR instrument at room temperature. Chemical shifts (δ) are expressed in ppm and were referenced against the solvent resonances at δ_H_ 7.26 and δ_C_ 77.23 ppm for CDCl_3_, and δ_H_ 4.87 and δ_C_ 49.15 ppm for CD_3_OD for ^1^H and ^13^C NMR respectively. HRMS were recorded on a Thermo Scientific Orbitrap Fusion spectrometer. The purity of compounds was checked using ^1^H NMR or thin layer chromatography (TLC) using pre-coated aluminium-backed plates (silica gel 60 F_254_, Merck) and compounds were visualised by UV radiation at 254 nm and then using an anisaldehyde spray reagent (1% *p*-anisaldehyde:2% H_2_SO_4_: 97% cold MeOH) followed by heating. Final purifications were done using preparative thin layer chromatography (Merck 818133) and gravity column chromatography that was carried out using a 2 cm diameter column, which were packed with silica gel (Merck Art. 9385) in selected solvent systems.

Dried leaves of *Vepris hemp* were ground to fine powder using a blender. The dried leaves 17.1 g were successively extracted, initially, using methylene chloride (CH_2_Cl_2_), and followed by methanol (CH_3_OH) solvents to yield 2.11 g and 1.63 g respectively. The methylene chloride extract was subjected to gravity column chromatography packed with a 1:1 blend of silica gel Merck 9385 and eluted using a step gradient initially starting with 100% hexane (250 mL), then 20% CH_2_Cl_2_ in hexane (250 mL), then 50% CH_2_Cl_2_ in hexane (250 mL), then 80% CH_2_Cl_2_ (250 mL), and 100% CH_2_Cl_2_ (250 mL), and finally 5% ethyl acetate (EtOAc) in CH_2_Cl_2_ (250 mL) was used. 10–15 mL, fractions were collected, and concentrated to dryness. The fractions were monitored using TLC and fractions with the same retention times were pooled. Fractions 7–9 gave lupeol, whereas fractions 14 – 17 gave arborinine (**2**) that we recently reported from *V. teva* (*Langat, Kami & Cheek, 2022*), and regularly reported from the genus *Vepris* (*Ombito, Chi & Wansi, 2021*). Fractions 27–33 gave kokasuganine (**1**), that we recently reported from *V. teva* (*Langat, Kami & Cheek, 2022*), and regularly reported from the genus *Vepris* (*Pusset et al., 1991; Ombito, Chi & Wansi, 2021*). Fractions 62–66 gave compound **8**, tecleanthine, which we previously reported from *V. verdoorniana* (*Atangana et al., 2017*), and previously from seven *Vepris* species (*Ombito, Chi & Wansi, 2021*). Fraction 67–69 gave compound **7**, evoxanthine which we previously reported from *V. verdoorniana* (*Atangana et al., 2017*), and previously from seven *Vepris* species (*Ombito, Chi & Wansi, 2021*). Fraction 71–72 gave 1,2,3,5-tetramethoxy-*N*-methylacridone (**4)**, which we previously reported from *V. gossweileri* (*Langat et al., 2021*), and *Vepris verdoorniana* (*Atangana et al., 2017*). Fraction 87–89 gave compound **3**, 1,2,3-trimethoxy-*N*-methylacridone, which was previously reported from *V. bilocularis* Engl.(*Ombito, Chi & Wansi, 2021*), whereas fraction 90–92 gave 1,3-dimethoxy-*N*-methylacridone (**5**) that is rampant in *Vepris* genus (*Ombito, Chi & Wansi, 2021*) and fraction 114–117 gave compound **6**, toddaliopsin A, which has been reported only once from, *V. bremekampii* (I.Verd.)Mziray (*Naidoo et al., 2005*).

The CH_3_OH extract was subjected to gravity column chromatography packed with a 1:1 blend of silica gel Merck 9385 and eluted using a step gradient initially starting with 100% CH_2_Cl_2_ (250 mL), then 1% CH_3_OH in CH_2_Cl_2_ (250 mL), then 2% CH_3_OH in CH_2_Cl_2_ (250 mL), then 5% CH_3_OH (250 mL), and 20% CH_3_OH in CH_2_Cl_2_ (250 mL).Fractions (10-15 mL) were collected, and concentrated to dryness. The fractions were monitored using TLC and fractions with the same retention times were pooled. Fractions 20–58 consisted of a mixture of compounds, which the ^1^H NMR spectrum showed to be similar to the compounds identified from the CH_2_Cl_2_ extract. From 67–69 we obtained isoplatydesmine (**11**), previously reported from *V. soyauxii* (Engl.)Mziray, *V. nobilis* (Delile)Mziray, *V. simplicifolia* (I.Verd.)Mziray and *V. tabouensis* (<u>Aubrév.</u> & <u>Pellegr.</u>)Mziray (*Ombito, Chi & Wansi, 2021*), whereas fraction 70 – 73 gave *N-*methylplaty-desminium ion (**9**) previously reported from *Araliopsis tabouensis* <u>Aubrév.</u> & <u>Pellegr.,</u> (now *Vepris*), *A. soyauxii* Engl., *Ruta graveolens* L. and *Skimmia japonica* Thunb. (*Ngadjui, Ayafor & Sondengam, 1988; Ombito, Chi & Wansi, 2021*), fraction 76–78 gave ribalinium ion (**10**) known to occur in *Balfourodendron riedelianum* (Engl.) Engl. and *Ruta chalepensis* L. (*Al-Majmaie, 2019*), and fraction 90 –94 gave the common, ferulic acid.

### Extinction risk assessment

The conservation assessment was made using the categories and criteria of *IUCN (2012)*. Threats were observed by the second authors directly in the field in Tanzania.

## RESULTS

### Chemistry

The structures of the compounds identified from the leaves *of V. hemp* were determined using extensive spectroscopic and spectrometric analysis, and the spectra of the known compounds compared to those previously reported (see table in supplementary materials). The compounds isolated from *V. hemp* were four quinoline alkaloids, kokusaginine (**1**) (*Pusset et al., 1991, Ombito, Chi & Wansi, 2021*), *N*-methylplaty-desminium ion (**9**) (*Ngadjui, Ayafor, Sondengam, 1988; Ombito, Chi & Wansi, 2021*), ribalinium ion (**10**) (*Al-Majmaie, 2019*), and isoplatydesmine (**11**) (*Ombito, Chi & Wansi, 2021*) (Fig. 1 & 2), and seven acridone alkaloids, arborinine (**2**) (*Langat, Kami & Cheek, 2022; Ombito, Chi & Wansi, 2021*), 1,2,3-trimethoxy-N-methylacridone (**3**) (*Ombito, Chi & Wansi, 2021*), 1,2,3,5-tetramethoxy-*N*-methylacridone (**4**) (*Atangana et al., 2017; Langat et al., 2021*), 1,3-dimethoxy-*N*-methylacridone (**5**) (*Ombito, Chi & Wansi, 2021*) and toddaliopsis A (**6**) (Naidoo et al., 2005), evoxanthine (**7**) (*Atangana et al., 2017; Ombito, Chi & Wansi, 2021*) and tecleanthine (**8**) (*Atangana et al., 2017, Ombito, Chi & Wansi, 2021*) (Fig. 1 & 2). In addition, lupeol and ferulic acid were isolated from this plant. Quinoline and acridone alkaloids are restricted to the *Rutaceae* family (*Basa & Tripathy, 1984; da Silva, Gottlieb & Ehrendorfer, 1988; Imbenzi et al., 2014; Michael, 2007; Michael, 2017*; *Ombito et al., 2020; Sackett, 2000*), confirming beyond reasonable doubt the placement of this material in the Rutaceae. Given the morphology (see introduction above and morphology below) placement within Rutaceae in Africa must be in the genus *Vepris*, where morphology and ecology suggest a close, possibly sister relationship with *Vepris uguenensis* Engl. (see morphology below). *Vepris uguenensis* has been studied for its chemistry (*Cheplogoi et al., 2008; Kiplimo, Islam & Koorbanally, 2012*). It has been recorded to contain flindersiamine, maculosidine, **ugenenazole, ugenenonamide, *N*-methyl-*N*-benzoyl-1-acetyltryptamide, methyl ugenenoate**, limonyl acetate, **ugenensene, ugenensone, niloticin, chisocheton A, kihadalactone A, tricoccin S**_**13**_ **acetate, 8α**,**11-elemodiol**, lupeol, **ugenenprenol**, and **syringaldehyde**. The 13 compounds in bold are unique to *V. uguenensis* within *Vepris* and notably have not been recorded by us (this paper) from *Vepris hemp*, giving strong chemotaxonomic support to the existing morphologically based species-level separation of *V. hemp* from *V. uguenensis*.

**Figure 1.**
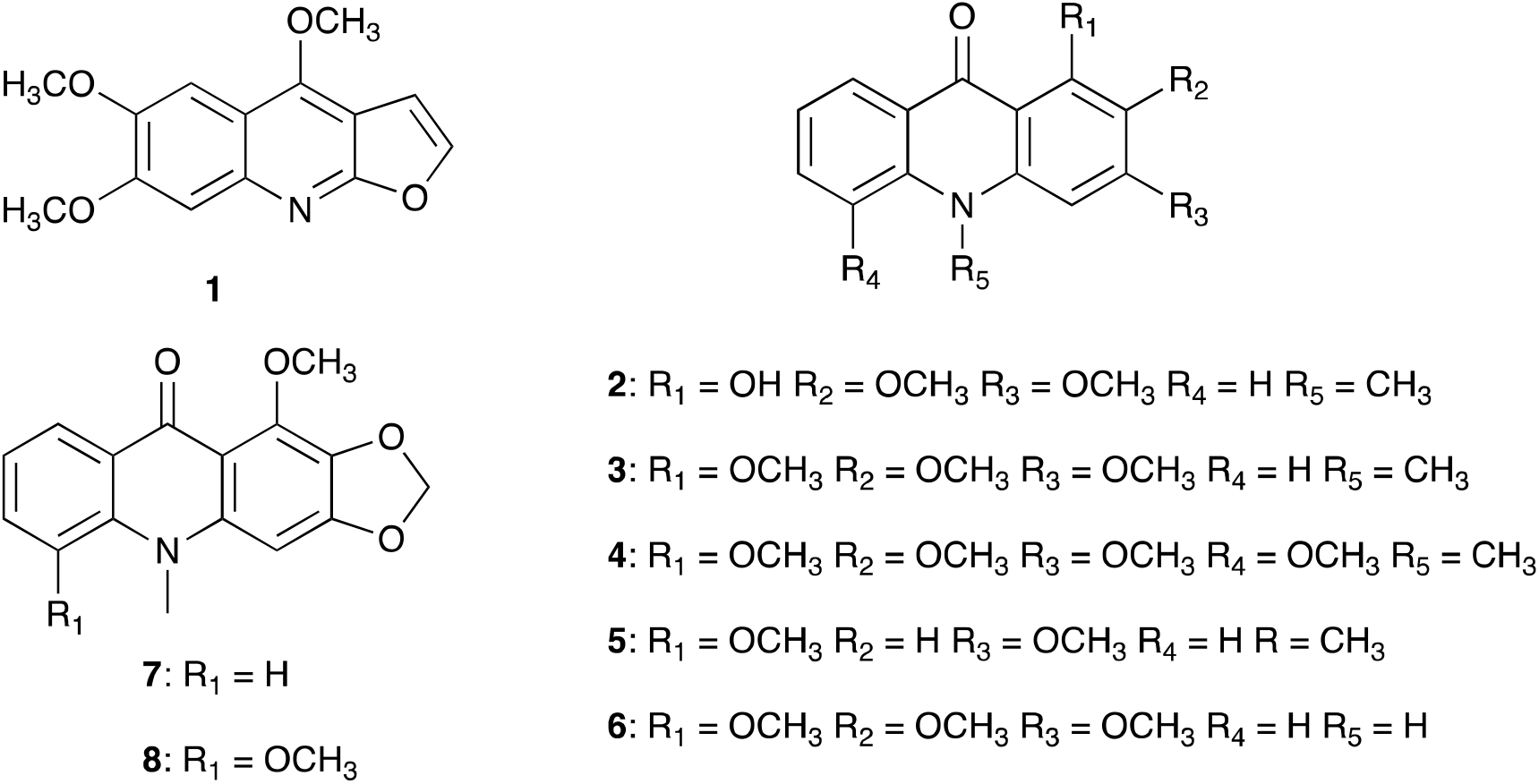
Structures of compounds identified from the methylene chloride of leaves of *Vepris hemp*

**Figure 2:**
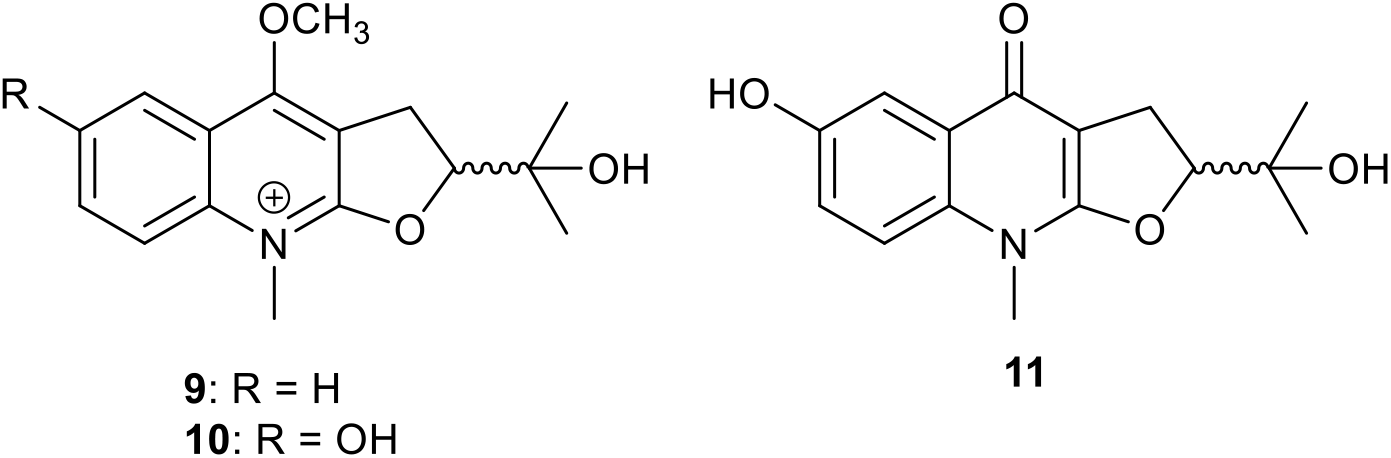
Structures of compounds identified from methanol extract of leaves of *Vepris hemp*

### Morphology

The morphology of *Vepris hemp* is unusual in the genus because of the high level of dense branching of the numerous slender leafy shoots, which results in a resemblance to a hedge plant (see Fig 2). Most other species of the genus are much more sparingly branched, see e.g. *Vepris teva* Cheek (*Langat, Kami & Cheek, 2022*). The very small, leathery leaflets and densely hairy stems and petioles of *Vepris hemp* suggest xerophytic adapations consistent with the habitat of deciduous woodland where insolation levels at the shrub layer in the dry season can be expected to be high due to lack of leaves in the forest canopy layer. Other species of *Vepris* are mainly confined to evergreen forest and perhaps for this reason do not show such xerophytic adaptions. Another exception which shows similar morphology, is *Vepris uguenensis* which atypically in the genus is recorded from grassland and savannah vegetation rather than forest (*Kokwaro, 1982*). We compiled a species identification key (see below) to the alternate, trifoliolate, sessile leafleted E African *Vepris* species reconstructed from *Kokwaro* (*1982*) by combining together species entries from his separate keys for the several genera united under *Vepris* since the work of *Mziray* (*1992*). Using this key, *Hemp* 7152 would be identified as *Vepris uguenensis*. However, the two species differ in numerous morphological characters (see diagnosis and Table 1). These are more than sufficient to warrant species recognition for *Hemp* 7152 as *Vepris hemp*. We updated the key by including data from *Vepris hemp*.

**Table 1.**
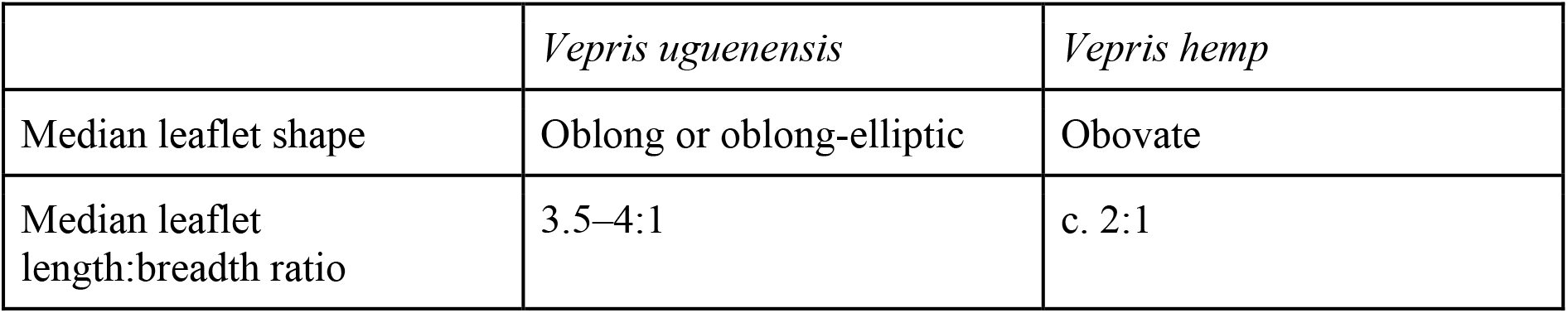
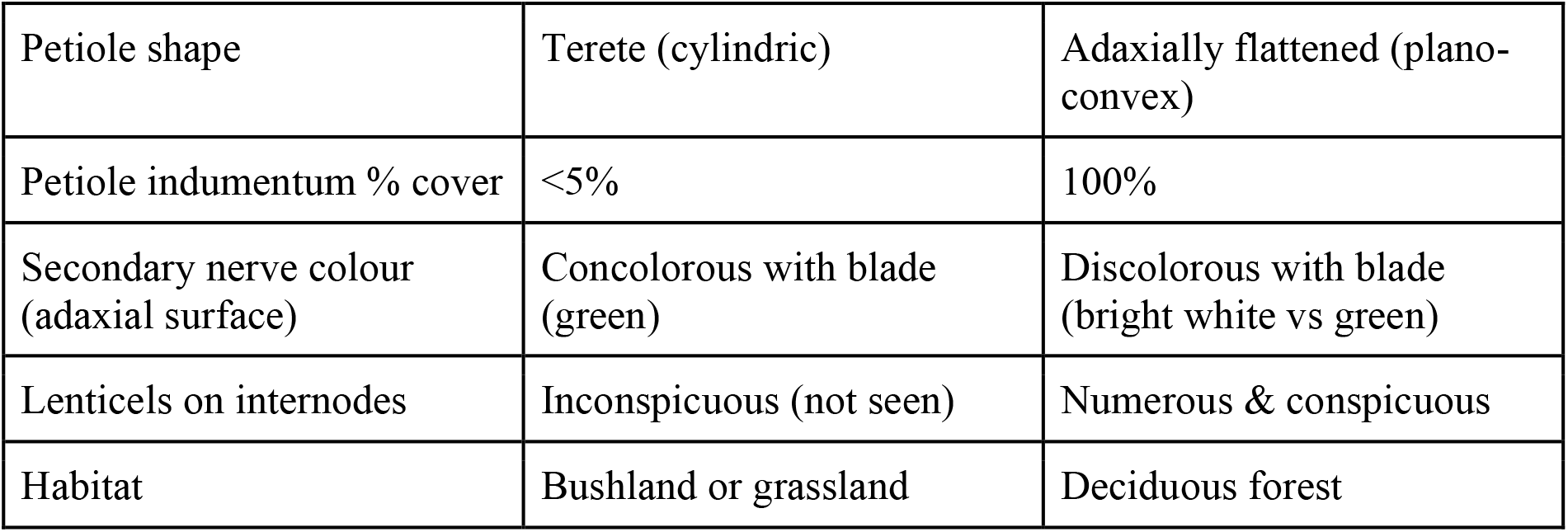
Characters separating *Vepris uguenensis* from *Vepris hemp*.

Since *Vepris uguenensis* occurs in the Pare Mts adjacent to the Usambara Mts, it is possible that the two are sister species, but molecular phylogenetic studies are needed to test this hypothesis.

**Key to the species of E. African (Uganda, Kenya, Tanzania) *Vepris* with trifoliolate, alternate, leaves and sessile leaflets**.

Based on data from *Kokwaro* (*1982*).

1. Leaves glabrous …………………………………………………………………… 2

1. Leaves hairy (minutely pubescent to villose) ……………………………………………………5

2. Leaflets obovate-cuneate (widest in the distal half), 4–6 cm long…***Vepris schliebenii***

2. Leaflets elliptic or elliptic-oblong (widest in the middle), 5–19 cm long ……………3

3. Fruit 4-lobed, ripening black ……………………………………… ***Vepris lanceolata***

3. Fruit unlobed, ripening red or orange ……………………………………………….4

4. Inflorescence-infructescence glabrous ………………………………… ***Vepris nobilis***

4. Inflorescence hairy ……………………………………………… ***Vepris grandifolia***

5. Leaflet bases auriculate; leaflets large 4–11 cm wide ……….….. ***Vepris samburensis***

5. Leaflet bases cuneate, rounded or truncate; leaflets, 0.8–7.5 cm wide ………….… 6

6. Deciduous; petiole winged ………………………………………. ***Vepris glomerata***

6. Evergreen; petiole grooved, not winged (except occasionally narrowly winged in *V. trichocarpa)*. …………………………………………………………………………………………… 77.

Longest leaflets on a specimen exceeding 6 cm long ……………………..… 8

7. Longest leaflets on a specimen less than 6 cm long …………………………………9

8. Inflorescence paniculate ……………………………..…………..***Vepris eggelingii***

8. Inflorescence racemose …………………………..………….….. ***Vepris trichocarpa***

9. Petiole terete (cylindrical); median leaflet shape oblong or oblong-elliptic; length:breadth ratio 3.5–4:1 …………………………………… ***Vepris uguenensis***

9. Petiole flattened adaxially (piano-convex); median leaflet shape obovate; length: breadth ratio, <2:1 ……………………………………………. ***Vepris hemp sp.nov***.

***Vepris hemp*** Cheek & Langat sp. nov. – Fig. 3.

**Fig. 1.**
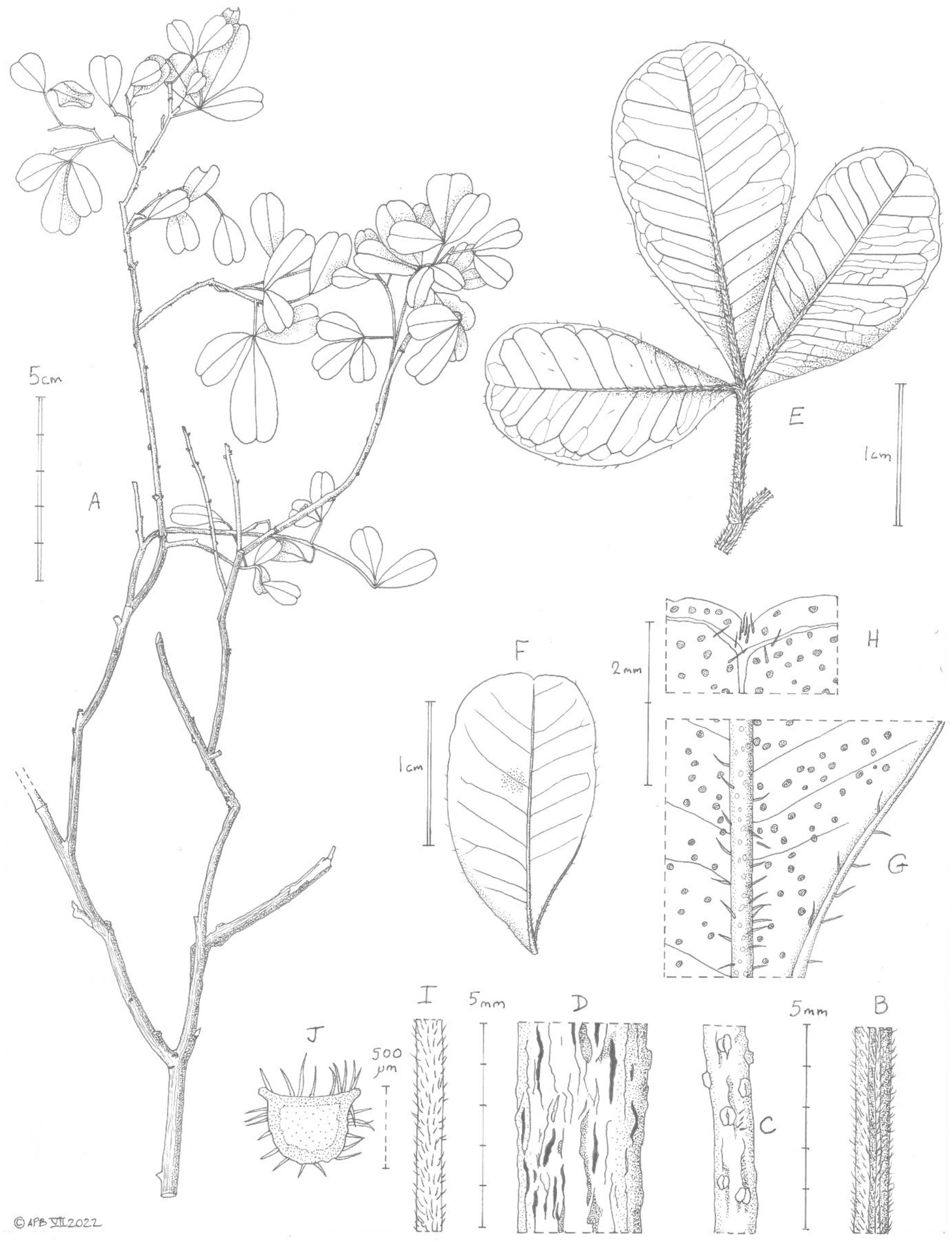
***Vepris hemp*** Cheek. (A) habit, leafy branch; (B) stem internode near apex: (C) stem four internodes from apex with lenticels: (D) major branch surface, lenticels absent; (E) leaf and stem, adaxial surface; (F) lateral leaflet, abaxial surface showing detail of surface glands; (G) detail of proximal part of (F) showing hairy midrib and margin, and conspicuous surface glands; (I) petiole, adaxial surface; (J) transverse section of petiole. Drawn by Andrew Brown from *Hemp* 7152.

Type: Tanzania, Tanga Region, West Usambara Mts, Mombo District, Mbale Escarpment, E 38,256579, -4,444915 S, 860m asl, relevé no. 2093, st. 9 Oct. 2018, *Hemp* 7152 (holotype K barcode K000593356; isotypes NHT, UBT).

Diagnosis: similar to *Vepris uguenensis* Engl. in the densely branched hairy stems, sessile, retuse leaflets wider in the distal than in the proximal halves, differing in the leaflets obovate, median leaflet length: breadth ratio c. 2:1 (vs oblong or oblong-elliptic, 3.5–4:1), petioles adaxially flattened (plano-convex), 100% hair covered (vs cylindrical (terete), <5% cover), lenticels on stems numerous & conspicuous (vs inconspicuous (not seen))

*Evergreen shrub, highly branched*, c. 2 m tall. *Principal stem* grey, terete with longitudinal blade-like ridges, 4–5 mm diam., c. 1.5 m above ground-level, glabrous, lacking lenticels. Leafy stems erect, terete, with 4–5 acute longitudinal ridges, 0.9–1.25 mm diam., internodes (1–)2–13 mm long, densely hairy, hairs covering about 50% of the surface of the second to fourth internodes (the first internode when young totally covered in hairs), hairs simple, patent, dull yellow or white, straight, tapering to an acute point, 0.25–0.3 mm long; lenticels scattered on the fourth and more distal internodes, white, raised, longitudinally elliptic, usually with a median longitudinal groove, 0.4–0.5 × 0.2 mm. *Leaves alternate, spirally arranged, trifoliolate*. Leaflets unequal, sessile, obovate, median leaflet longer than the laterals, symmetrical, 13.5–28 × 6–14 mm; lateral leaflets strongly asymmetric at base, 9.5–22 × 5.5–11 mm, apex retuse, the sinus 0.4–0.5 mm deep, c. 2 mm wide, with conxex sides; base acute, thinly coriaceous, adaxial surface glossy green, midrib, secondary and tertiary nerves bright white, conspicuous and raised, abaxial surface pale matt green, midrib and proximal third of secondary nerves white, remaining nerves concolorous with blade, inconspicuous. Secondary nerves 7–8(–9) on each side of the midrib, arising at 70–80 degrees from the midrib, straight, near the margin curving or angled upwards sharply, and uniting with the nerve above, forming an angular to looping infra-marginal nerve, 0.8–1.2 mm from the margin, usually with a parallel, weaker infra-marginal nerve 0.05–0.1 mm from the margin; tertiary nerves parallel to the secondaries but highly branched; gland dots translucent, colourless in transmitted light, conspicuous; on adaxial surface concolorous, inconspicuous; on abaxial surface cblack or yellow-brown, raised, 9–18 per mm^2^; indumentum as the stem, dense along abaxial midrib, mixed with raised glandular areas, terminus of midrib with a tuft of hairs, adaxial midrib glabrescent at length, margins sparsely hairy. Petiole terete, 4–10 × 0.5 mm, densely hairy as the stem.

### Distribution

Tanzania, Tanga Region, West Usambara Mts, Mombo District, Mbale Escarpment.

### Ecology

*Vepris hemp* appears to be restricted to a rare and almost extinct type of dry deciduous forest in the Usambara Mts of Tanzania. Here we characterise the vegetation at the site of the type locality of *Vepris hemp:*

***Commiphora* dry deciduous forest** on a steep (inclination 35°), rocky (boulders 0.3–2m covering 15% of the plot), ne exposed slope with about 80 species on 0.1 ha. pH was 6.9 (H_2_0) and 6.5 (CaCl_2_). The tree layer had a cover of 60% and a height of 18 m. Dominant tree species were *Commiphora baluensis, Sterculia africana, Erythrina sacleuxii, Euphorbia quinquecostata* and *E. bussei*. The dense shrub layer of 60% cover and 4 m height was very diverse including species such as *Acalypha fruticosa* var. *eglandulosa, Combretum exalatum, Abrus schimperi* ssp. *africanus, Dombeya taylorii* and *Pentas parvifolia*. In the herb layer *Adiantum incisum, Panicum deustum, Cyperus glaucophyllus, Xerophyta spekei* and *Barleria submollis* were prominent. The new *Vepris* species occurred with low frequency and a cover of less than 1% in the shrub layer of this vegetation plot. Its height was about 2 m. The small forest patch of about only 1.5 ha was quite undisturbed since it was located in a very steep and difficult to access part of the escarpment. However, it was completely surrounded by relicts of destroyed former dry forests and by bushlands heavily degraded due to grazing, and due to timber and firewood extraction.

### Local names and uses

None are recorded.

### Etymology

*Vepris hemp* is named for Dr Andreas Hemp, collector of the type specimen. Based at University of Bayreuth, Germany, he has conducted plant ecology and biodiversity research in Tanzania since 1989 and since 2010 has led the project “Kilimanjaro ecosystems under global change” and “The role of nature for human well-being in the Kilimanjaro Social-Ecological System” (abbreviated to KiLi and Kili-SES, see https://www.kilimanjaro.biozentrum.uni-wuerzburg.de/ and https://kili-ses.senckenberg.de/) together with Markus Fischer (Bern), Katrin Böhning-Gaese and Claudia Hemp (both Würzburg). Building research capacity in Tanzania and engagement with local communities is an important aspect of his work.

### Conservation

Known from a single location on the slopes of the West Usambara Mts of Tanzania, where it grows in a rare surviving fragment of an unusual dry forest vegetation type (see Ecology), *Vepris hemp* is here assessed as Critically Endangered since only a single site is known, area of occupation is calculated as 4 km^2^ using the preferred IUCN grid cell size and extent of occurrence as the same. The forest fragment is mainly under pressure of clearance for timber, charcoal and grazing and is unprotected apart from its steep slope and the difficulty in accessing it, which may be lost in future. This justifies an extinction risk assessment of CR B1ab(iii) + B2ab(iii) using the *IUCN* (*2012*) standard. Numerous other plant species are globally restricted to the Usambara Mts and are also threatened with extinction due to human pressures e.g. *Vepris amaniensis* (Engl.) Mziray (*Cheek & Luke, 2022*) and *Cola lukei* Cheek (*Cheek, 2002*).

### Notes

Efforts should be made to recollect *Vepris hemp* in flower and in fruit in order to complete its taxonomic characterisation, and also to obtain seed to attempt to cultivate it, to reduce the high risk of its extinction. We would expect the flowers to have four sepals and four petals as usual in *Vepris*, and likely 4 or 8 stamens in the male flowers, with in the female, a single entire, bilocular pistil followed by an entire, 1-seeded berry, as in *Vepris ugenensis*.

This discovery takes to 22 the number of formally described species of *Vepris* in the FTEA area (Uganda, Kenya, Tanzania), with 19 recorded by *Kokwaro* (*1982*) and a further three published recently (*Cheek & Luke, 2022*).

## CONCLUSIONS

The case of *Vepris hemp*, seemingly on the edge of extinction, illustrates the importance of discovering previously unknown species and bringing them to light by publishing them as new to science, if possible even when flowers and fruit are unknown, as here. Formal naming facilitates acceptance by IUCN of extinction risk assessments (Red Listing) which increases the possibiity of resources being allocated to support conservation. Over the last 15 years the number of new species of vascular plant published has remained more or less constant at about 2000 species per annum (*Cheek et al., 2020*) adding to the the c. 369,000 species already accepted, although this number is disputed (*Nic Lughadha et al., 2016*). Many of these species have not been discovered previously because, like *Vepris hemp*, they appear severely range-restricted, and have been hidden until their location has been studied botanically, in this case thanks to the network of plots established by several DFG-funded projects including the KiLi-Project to monitor vegetation change (*Hemp, 2012*). This range-restriction of new species makes them almost automatically threatened because threats are now so ubiquitous in tropical Africa. However, only 7.2% of vascular plant species have been assessed and included on the Red List using the *IUCN* (*2012*) standard (*Bachman et al., 2019*).

Extinctions of plant species due to habitat clearance are increasing, e.g., in the vicinity of *Vepris hemp* in the Usambara Mts of Tanzania, *Cynometra longipedicellata* Harms may well now be extinct at its sole locality (it is always difficult to be 100% certain that extinction has occurred), the Amani-Sigi Nature Reserve (*Gereau et al., 2016*). Also in Tanzania, *Kihansia lovettii* Cheek (Triuridaceae, *Cheek, 2004*) at the Kihansi dam site, has not been seen since it was first collected despite targeted searches. Global plant species extinctions are being recorded across Africa, e.g. in Cameroon, species of non-photosynthetic mycoheterotroph *Oxygyne triandra* Schltr. and *Afrothismia pachyantha* Schltr. (Thismiaceae, *Cheek & Williams, 1999; Cheek et al., 2018; Cheek, Etuge & Williams, 2019*) and Podostemaceae such as *Inversodicrea bosii* C.Cusset (*Cheek et al., 2017*), while in Gabon the spectacular *Pseudohydrosme buettneri* Engl. is also thought to be extinct having not been seen for over 100 years despite searches for this genus (Araceae, *Cheek, Tchiengué & van der Burgt., 2021*). Examples of species apparently becoming extinct even before they are formally known to science are on the rise, e.g. *Saxicolella deniseae* Cheek in Guinea (Podostemaceae, *Cheek et al., 2022*), *Vepris bali* Cheek in Cameroon (Rutaceae, *Cheek, Gosline & Onana, 2018*), *Pseudohydrosme bogneri* Cheek & Moxon-Holt in Gabon (Aracae, *Moxon-Holt & Cheek, 2021*) and in the Uluguru Mts of Tanzania *Vepris sp. A* of FTEA which is still formally not named (Rutaceae, *Cheek & Luke, 2022*). Each of these global extinctions represents the loss of opportunities for humanity. In the case of *Vepris*, not only are many of the species used for traditional medicine, but research has shown that the different species each produce an array of bioactive compounds in the alkaloid class, with potential applications, e.g. with potent levels of antimicrobial activity, especially in combination and with compounds new to science being discovered frequently as more species are investigated (*Langat, Kami & Cheek, 2022)*.

In Tanzania, on the whole, natural habitat is relatively well covered by a well-planned network of protected areas, but nevertheless natural habitat at some sites with species of high value for conservation has all but disappeared or is at risk of disappearing as for the site for *Vepris hemp* (see above). Further investment in prioritising the highest priority areas for plant conservation as Tropical Important Plant Areas (TIPAs, using the revised IPA criteria set out in Darbyshire *et al*. (2017)) as is in progress in countries such as Guinea, Cameroon, Uganda, Mozambique and Ethiopia might be extended elsewhere to reduce further the risk of future global extinctions of range-restricted endemic species such as *Vepris hemp*.

## Supporting information

Supplemental morphological data

## ACKNOWLEDGEMENTS

Andreas Hemp gratefully acknowledges the financial support granted by the Deutsche Forschungsgemeinschaft and the research permit granted by COSTECH and TAWIRI. Dr Neduvoto Mollel, Director of NHT, is sincerely thanked for allowing access to the NHT Herbarium. Martin Cheek thanks Janis Shillito for typing and Rosemary Lomer for data entry.

